# Increased substrate complexity drives re-diversification and functional reorganization in simplified methanogenic consortia

**DOI:** 10.64898/2026.09.01.748264

**Authors:** Raphaëlle Péguilhan, Ginevra Giangeri, Marlene Mark Jensen, Irini Angelidaki

## Abstract

Anaerobic digestion is a sustainable process for methane production that relies on complex microbial networks. While simplified enriched consortia offer a promising strategy to improve process control, excessive simplification can disrupt key functions and microbial partnerships, reducing community resilience. In this study, we investigated whether simplified methanogenic communities could re-diversify and maintain methane production when exposed to more complex substrates, namely butyrate and glucose. We also evaluated the effect of vitamin and amino acid supplementation on sustaining key methanogens and beneficial microbial partners. Three methanogenic communities were monitored over three months for methane production and microbial diversity while receiving butyrate and/or glucose, with different vitamin or amino acid supplements. Exposure to more complex substrates successfully restored the diversity of acidogenic and acetogenic populations, even after prolonged feeding with simple substrates, highlighting both the resilience of the simplified communities and the ecological importance of low-abundance taxa. However, the transition reduced process stability and methane production, likely due to substrate overloading. The results further suggest that substrate complexification should be introduced stepwise, promoting acetogenesis before acidogenesis. This fundamental study brings new light on which factors must be considered in the long-term goal of designing tailored-made consortia for anaerobic digestion.

**One sentence summary:** Increasing substrate complexity re-diversified simplified methanogenic consortia, revealing the importance of low-abundance taxa and functional resilience in anaerobic digestion communities.

## Introduction

Anaerobic digestion (AD) is a sustainable biotechnological process that converts waste into a valuable product, such as biomethane. This process relies on key anaerobic steps, namely hydrolysis, acidogenesis, acetogenesis and methanogenesis, to break down highly complex substrate, such as feedstock, into methane. The efficiency of this process depends heavily on the balance between these different steps, which are carried out by a complex microbial network (Adekunle and Okolie 2015). The microorganisms responsible for the substrate degradation at each step of the AD vary greatly in terms of physiology, nutritional requirements, growth kinetics and sensitivity to the environment. Consequently, it can be difficult to maintain a balance within these microbial communities between acid forming and methane forming microorganisms, which often leads to AD with low methane yield and an accumulation of volatile fatty acids (VFAs) (Adekunle and Okolie 2015, Xu F *et al*. 2018). To tackle the challenges related to AD and overly complex microbial communities, new ambitious strategies have emerged such as the design of simplified or tailor-made microbial consortia (Giangeri, Campanaro *et al*. 2025). Simplified microbial consortia can be designed through the synthetic assembly (bottom-up approach) of selected and already characterized microorganisms, or results from targeted enrichment (top-down approach) aimed at reducing the diversity of an inoculum and retaining only a simplified community geared towards a specific process (Lyu *et al*. 2024, Jourdain and Gu 2025). While such enriched simplified community can be promising to improve the production or degradation of specific compounds, it comes with several limitations (Agler *et al*. 2014, Gilmore *et al*. 2019, Tukanghan *et al*. 2021, Hu *et al*. 2022). It is primordial to maintain sufficient diversity within a simplified community in order to avoid any unexpected impact on its functioning and loss of beneficial interactions, as illustrated by the bottleneck effect (Abel *et al*. 2015, J Cira, Pearce, and Quake 2018). Low-abundance taxa can play a key role within the microbial community and help to maintain its resilience when new substrates or stressors are introduced (Shade *et al*. 2014).

With regard to methanogenesis, the final and critical step of AD, this can be major limiting factor in the process, as methanogens heavily rely on other microbial partners to degrade complex substrate into acetate, H2 and CO2. Some studies therefore focus on enriched communities, fed directly with methanogenic substrates, to investigate the process more efficiently and reduce the risk of unexpected imbalances or rate limitations caused by inefficient breakdown of more complex substrates into acetate or H2:CO2 (Chang *et al*. 2025, Giangeri, Tsapekos *et al*. 2025). However, as previously mentioned, the direct use of methanogenic substrates can lead to highly simplified communities that will lose their resilience, for example, when exposed to more complex substrates, and will lose potentially syntrophic or mutualistic microbial interactions that promote methanogenesis. These beneficial interactions can take the form of an exchange of essential nutrients, such as vitamins or amino acids, between syntrophic or mutualistic microbial partners (Sieber, McInerney, and Gunsalus 2012, Hubalek *et al*. 2017). Furthermore, methanogens have other limitations, such as slow growth compared with acidogenic or acetogenic bacteria and high vulnerability to various stress factors (Chen, Cheng, and Creamer 2008). Members of *Methanosarcina* genus are considered to be more robust to stress (e.g., low pH, high ammonium and volatile fatty acid [VFA] concentrations) than other methanogens. In addition, they are metabolically versatile archaea that can perform hydrogenotrophic, acetoclastic and methylotrophic methanogenesis, allowing potentially more flexibility in the system (De Vrieze et al. 2012). Therefore, *Methanosarcina* emerges as a suitable candidate to target when enriching microbial consortia to optimize methane production.

The main aim of this study was to evaluate if a simplified methanogenic community exposed to a more complex substrate will be able to reorganize and maintain methane production. The hypothesis is that low-abundance or dormant taxa that remained in the simplified community will be stimulated by the new substrate and help the recovery of the AD process. A second objective was to investigate if supplementations in vitamins and amino acids can help maintain the methanogens, such as *Methanosarcina* spp., during the substrate disturbance.

The hypothesis is that *Methanosarcina* and potential identified partners can be maintained or enriched by complementing their predicted auxotrophies. In this aim, simplified AD consortia, previously optimized for methane production and fed with acetate and H2:CO2, were used as model AD communities (Giangeri, Tsapekos *et al*. 2025). These communities were exposed to more complex substrates, e.g., glucose to stimulate acidogenesis and butyrate to stimulate acetogenesis, to re-diversify the community. Moreover, to support methanogenesis during this disturbance, the communities were supplemented with amino acids and vitamins intended to support the growth of *Methanosarcina* and its potential microbial partners. This fundamental study addressed the long-term goal of constructing synthetic microbial consortia by deepening our understanding of AD microbial community dynamics and the role of low-abundance taxa for process resilience.

## Materials and Methods

### Experimental setup for substrate enrichment

Three microbial communities originating from Hashøj Biogas plant (Denmark), which had been enriched for methane production with acetate or H2:CO2 over five generations, were selected from the previous work of (Giangeri, Tsapekos *et al*. 2025): Ac (acetate-fed), H2:CO2

PANI (H2:CO2-fed with cylindrical plastic materials coated with polyaniline [PANI]) and, H2:CO2 mag (H2:CO2-fed containing magnetite). These three cultures were selected according to their methane production and high abundances of *Methanosarcina* spp., as the main methanogen. They were used as inoculum for this study and renamed A, P and M, representing A for acetate, and the P and M represented inocula enriched with H2:CO2 with PANI and magnetite respectively.

As controls, the three inocula were kept under the same conditions as previously described in (Giangeri, Tsapekos *et al*. 2025): same substrate and conductive materials. Each control was cultured in a 250 mL serum bottle containing 80 mL of medium and 20 mL of initial inoculum. The core medium contained: basal anaerobic (BA) medium with 0.5 g/L yeast extract (Yeast Extract, powder, Ultrapure, Thermo Scientific), 1 mL/100 mL vitamins solution x10 and 0.25 g/L Na2S (Angelidaki and Sanders 2004) (see **Supplementary Material and Methods**). The controls were fed with either a H2:CO2 (4:1) gas mixture from gas cylinders (Air Liquide, Danmark A/S, Paris, France) supplemented every 3-4 days, or 2 g/L of sodium acetate (Sodium Acetate ≥99%, ACS ReagentPlus, Sigma-Aldrich) supplemented every 10 days. Conductive materials (PANI and magnetite) were not parameters examined in this study but were kept under H2:CO2-fed conditions to maintain the original cultivation medium and preserve the original community structure of the inocula.

Either 10 g/L of PANI or 10 g/L of magnetite (Iron(II, III) oxide powder 95 %, <5 μm, Sigma-Aldrich) was added.

Enriched conditions were done in duplicates in 120 mL serum bottles containing 40 mL of medium and 10 mL of inoculum. The medium was identical to the control, except for the substrate and amino acids and vitamins supplementation as described in Table 1. To assess the effects of higher substrate complexity than acetate or H2:CO2, butyrate and glucose were used, either individually or in combination: 2 g/L glucose (D-Glucose anhydrous, VWR Chemicals), 2 g/L sodium butyrate (Sodium butyrate 98%, Sigma-Aldrich) or 2 g/L 50:50 glucose and sodium butyrate (**Table 1**).

**Table 1:**
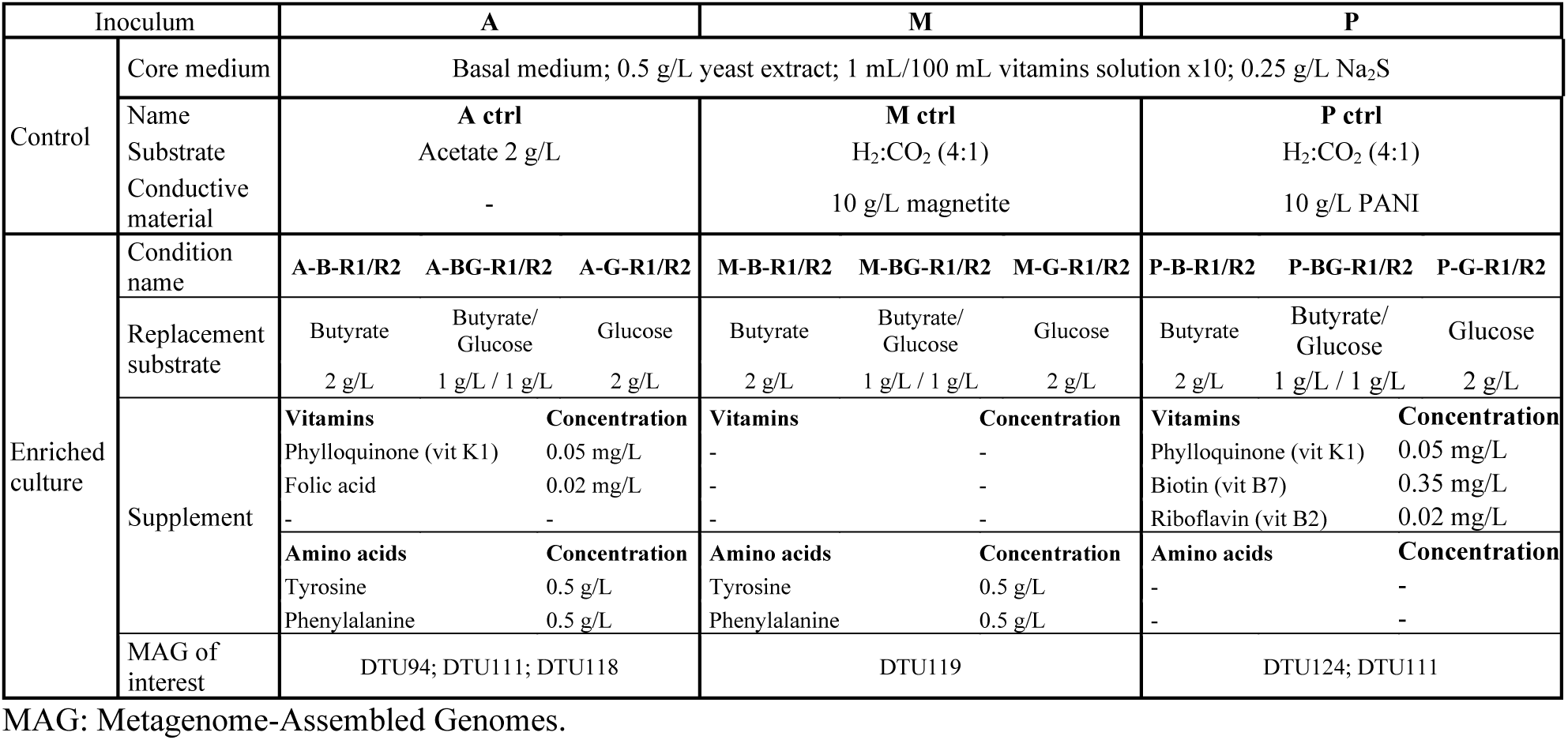
Medium and substrates used in controls and enriched cultures.

Moreover, to support the growth of *Methanosarcina* spp. and its potential microbial partners during the substrate change, selected amino acids and vitamins were included in the experiments as described in Table 1: 0.5 g/L phenylalanine (L-Phenylalanine 98.5-101.0%, BioXtra, Sigma-Aldrich), 0.5 g/L tyrosine (L-Tyrosine 99.0-101.0%, BioXtra, Sigma-Aldrich), 0.35 mg/L (0.36 mg/L final, in addition of the x10 vitamin solution) Biotin (Biotin ≥99%, BioReagent, Sigma-Aldrich), 0.05 mg/L Phylloquinone (Vitamin K1 ready Made Solution 5mg/mL, Sigma-Aldrich), 0.02 mg/L (0.07 mg/L final) Riboflavin ((-)-Riboflavin ≥98%, BioReagent, Sigma-Aldrich) and, 0.02 mg/L (0.04 mg/L final) Folic acid (Folic acid ≥97%, BioReagent, Sigma-Aldrich) (Table 1).

For the first two feedings, the enriched conditions related to inoculum A were fed with decreasing amounts of acetate to help the transition of the microbial community to butyrate or glucose: 1 g/L at 10 days and 0.5 g/L at 20 days. In the same way, enriched conditions for inocula P and M were fed a 4:1 H2:CO2 ratio every 3-4 days for the first three feedings. Then, the enrichment cultures were fed every 10-11 days with either butyrate or glucose alone. The cultures were reinoculated every 30 days in fresh medium and maintained for three generations. Enriched cultures were named based on the inoculum, the new substrate (B: butyrate; G: glucose; BG: Butyrate/Glucose) and the replicate number (R1 or R2) as follows: [A; M; P]-[B; G; BG]-[R1; R2].

### Gas and liquid phases analysis

The gas composition of the headspace in CH4, CO2 and H2 was monitored before and after each feeding using a GC-TRACE 1310, (Thermo Fisher Scientific, Waltham, MA, USA) Gas Chromatographer (GC), as reported in (Goonesekera *et al*. 2024). The VFAs concentration was monitored every 10 days before and after feedings during the third generation, when the enriched cultures were more stabilized. Liquid samples of 1.5 mL were acidified with 45 µL of H2O4S 10% (v:v) and centrifugated. Then, the supernatant was filter on 0.22 µm membrane (EconoFilter PTFE 13mm, Agilent Technologies, CA, USA) and 1 mL was stored at -20 °C until analysis. Glucose, butyrate, iso-butyrate, acetic acid, propionic acid, valeric acid, iso-valeric acid, caproic acid, and ethanol concentrations were monitored using a Nexera XR HPLC (Shimadzu Corporation, Kyoto, Japan) as reported in (Vayena *et al*. 2024) (**Supplementary data 1**).

### DNA extraction, amplification and sequencing

Volumes of 10-15 mL were taken from the three control bottles two months before the start of the enrichment to check the new composition of the microbial communities after reactivation, and then from each bottle every ∼15 days during the 3-month enrichment for DNA extraction and stored at -20°C until further processing. After 10 min centrifugation, DNA from pellets was extracted with the DNeasy PowerSoil Pro kit (Qiagen, Hilden, Germany) and quantified by fluorescence using the Qubit 4 Fluorometer (Invitrogen™, Thermo Fisher Scientific, USA) (**Supplementary data 2**). Inoculum A had previously been sequenced for shotgun metagenomics for other purposes than this study and was deposited under NCBI BioProject PRJNA1221661, sample SAMN46749652 (acetateNOPANI). Shotgun metagenomics for the two P and M inocula was done by DTU National Food Institute sequencing service (Lyngby, Denmark) using an Illumina NextSeq sequencing platform with a depth of 40M in duplicates for investigation of the inoculum before to plan the enrichment experiment. Finally, concerning the final point for enriched samples, shotgun sequencing was done by the company Majorbio Shanghai Meiji Biomedical Technology Co. LTD (Shanghai, China) using an Illumina NovaSeq 6000 platform (paired-end 2×150) to obtain 20M reads in the final raw data. Raw shotgun data have been deposited on the European Nucleotide Archive (ENA) at EMBL’s European Bioinformatics Institute (EBI) under the project accession PRJEB98414, samples SAMEA120801592 to SAMEA120801598. For metabarcoding, amplification and sequencing were done by Majorbio. The V4 region of the 16S rRNA gene was amplified by PCR (GeneAmp^®^ 9700 PCR system, Applied Biosystems, Thermo Fisher Scientific, USA) from total genomic DNA using the universal primers from the Earth Microbiome project, 515F 5’-GTGYCAGCMGCCGCGGTAA-3’, 806R 5’-GGACTACNVGGGTWTCTAAT-3’ (Apprill *et al*. 2015, Parada, Needham, and Fuhrman 2016). Amplicons were sequenced on the Illumina MiSeq platform using 2×250 bp paired-end. Demultiplexed raw data are available through the ENA under the project accession PRJEB98414, samples SAMEA120799023 to SAMEA120799148.

### Metabarcoding data processing

The 7,614,724 reads were processed with FROGS pipeline (v 5.0.2) (Escudié *et al*. 2018, Bernard *et al*. 2021) using DADA2 packages (v 1.22.0) (Callahan *et al*. 2016) to filter merged reads and to obtain amplicon sequence variants (ASVs) following FROGS guidelines. Only ASVs representing at least 5×10^-5^% of the total abundance were kept. The ASVs were affiliated with FROGS by BLAST and RDP assignations using SILVA v138.2 (Quast *et al*. 2013) as the reference database. Finally, a total of 6,787,069 sequences from all samples corresponding to 642 ASVs were kept for the analysis after all filtration steps (**Supplementary data 3**).

### Shotgun metagenomics data processing

An average of 20M-100M reads per sample were obtained from shotgun sequencing. Preprocessing of the data consisted of (i) FastQC (v 0.12.1) (Andrews 2010) to control raw data quality, (ii) Trimmomatic (v 0.39) (Bolger, Lohse, and Usadel 2014) to filter and trim erroneous read Trimmomatic was used with a sliding window of 4:15, a minimum read length of 65 bp and the leading and trailing parameters with a quality threshold at 30. This step removed between 2% to 16% of the raw reads. The trimmed reads were then co-assembled using MEGAHIT (v 1.2.9) (Li *et al*. 2015) with *meta-sensitive* option and a minimum contig length of 500 pb. The binning of the 164,624 obtained contigs (max length 834,835 bp, N50 15,568 bp) was executed with metaWRAP pipeline (Uritskiy, Diruggiero, and Taylor 2018) using CONCOCT (Alneberg *et al*. 2014), MaxBin2 (Wu, Simmons, and Singer 2016), metabat1 (Kang *et al*. 2015) and metabat2 (Kang *et al*. 2019) as binning tools and the *bin_refinement* and *reassemble_bins* modules. The 90 MAGs obtained were pooled with the 152 MAGs from (Giangeri, Tsapekos *et al*. 2025) and dereplication was done using dRep v 3.5.0 (Olm *et al*. 2017). Trimmed reads were mapped to the 196 dereplicated MAGs, using bowtie2 v 2.5.4 (Langmead and Salzberg 2012) with *no-mixed* and *no-discordant* options and the relative abundance of each MAGs was estimated with coverM v 0.7.0 (Aroney *et al*. 2025). The taxonomy of the 44 new MAGs was estimated with gtdbtk v 2.4.0 using the classify_wf module (database last access: Oct 2025) (Chaumeil *et al*. 2022, Parks *et al*. 2022) (**Supplementary data 4**). The 196 MAGs can be recovered from the ENA project PRJEB98414.

### Vitamin and amino acid auxotrophy screening

The data related to MAGs present in the inocula (i.e., DNA sequences, gene content, taxonomy and relative abundances) were retrieved from the study of (Giangeri, Tsapekos *et al*. 2025), NCBI BioProject PRJNA1063026. The main MAGs affiliated with *Methanosarcina* in the three inocula were manually refined, as described in the **Supplementary Materials and Methods**. COG annotations (Galperin *et al*. 2025), KEGG annotations (Kanehisa and Goto 2000, Kanehisa *et al*. 2025) and KEGG module (Kanehisa *et al*. 2017) completeness were inferred with Anvi’o 8 platform (Eren *et al*. 2020) using the programs *anvi-run-ncbi-cogs*, *anvi-run-kegg-kofams*, *anvi-export-functions*, and *anvi-estimate-metabolism* (Veseli *et al*. 2025). A KEGG module was considered as ‘complete/present’ if its completeness was ≥ 0.75. The pathwise completeness estimation strategy was chosen from the output for further analysis (**Supplementary data 5**).

Auxotrophies for vitamins and amino acids biosynthesis and carbon source degradation were screened using either KEGG module completeness or using enzyme hit (from module completeness analysis), COG genes and KEGG gene ID (KOfam database) when no KEGG module was available (**Supplementary data 5**). For amino acids, analysis by the GapMind web tool (Price, Deutschbauer, and Arkin 2020) was also used to confirm the KEGG predictions. In cases of contradiction, high-confidence KEGG module results were retained.

### Quantification of targeted MAGs by real-time PCR

The three MAGs DTU94, DTU119 and DTU124 affiliated with *Negativicutes* sp., Eubacteriales sp. and *Advenella* sp., and total bacteria were quantified by qPCR using primers targeting the 16S rRNA gene previously described in the literature (**Supplementary data 6**). Measurements were performed at T2 (Day 28), T4 (Day 57) and T6 (Day 88). The assay was performed in 20 µL of reaction mix containing 10 µM of forward and reverse primers (50:50 ratio with 2 reverse primers), 10 µL of FastStart Essential DNA Green Master (Roche, Basel, Switzerland), and 5 µL of DNA extract (2 ng/µL). The qPCR amplifications were performed on a LightCycler^®^ 96 instrument (Roche, Basel, Switzerland). All samples were measured in duplicate. The amplification parameters were as follows: 94-95 °C/2-10 min, followed by 39-45 cycles of [94-95 °C/30 s; annealing temp primer/30 s-1 min; 68-72 °C/1-2 min] (**Supplementary data 6**). The reference curves were generated using known concentrations of *Selenomonas ruminantium subsp. Lactilytica* DSM2872, *Bacillus licheniformis* DSM13, and *Advenella faeciporci* DSM104993 genomic DNA (Leibniz Institute DSMZ, Braunschweig, Germany).

### Normalization, statistics and visualization

RStudio (v 2024.12.1, R environment v 4.4.0) (R Core Team (2019), n.d.) with ‘vegan’, ‘ggdendro’, ‘ggsignif’, ‘compositions’ and ‘zCompositions’ packages were used for graphical representations, normalization and statistics on metabarcoding and process data. BioRender.com was used for the methanogenesis schematic diagram. ASV abundance data were normalized either by Total Sum Scaling (TSS) to calculate alpha-diversity indexes and obtain relative abundance or using centered log-ratio (CLR)-transformation as recommended by (Gloor *et al*. 2017) for Non-metric Multidimensional Scaling (NMDS) analysis.

## Results

### Process performances of the enrichment procedure

In the three control conditions, CH4 production continued at a relatively constant rate, while total H2 and CO2 were consumed within 2-3 days and acetate within 10 days (**Figure 1; Supplementary data 1**). The theoretical maximum yields of methanogenic substrate to CH4 produced (gCOD/gCOD) reached up to 54%, 14% and 74% over one generation of cultures in A ctrl, M ctrl and P ctrl, respectively (**Supplementary data 1**).

**Figure 1:**
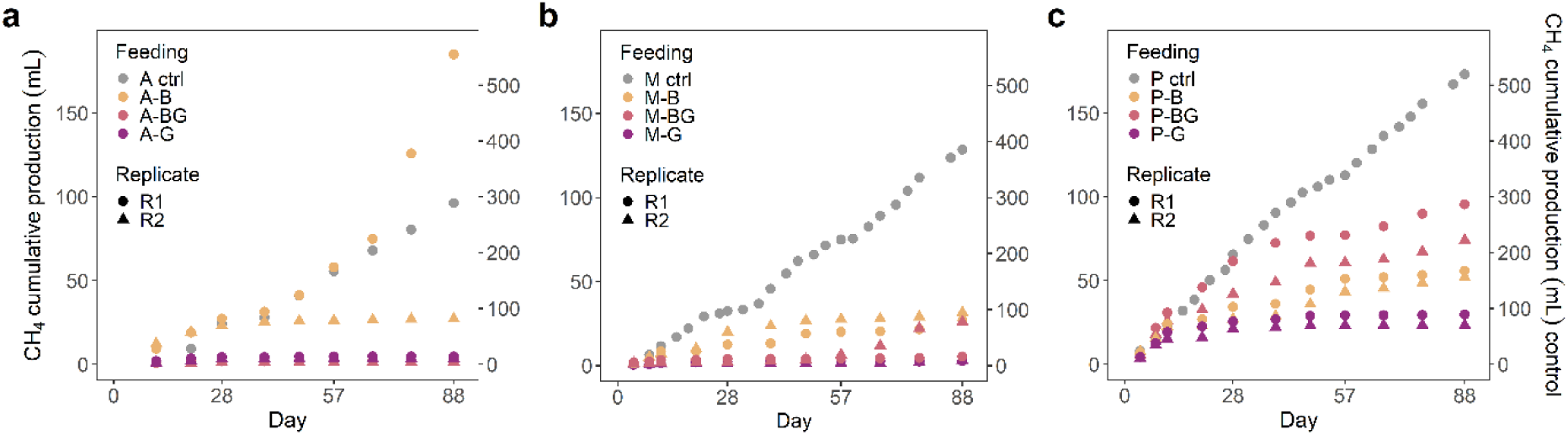
Cumulative methane production over three months for **a)** A, **b)** M, and **c)** P inocula. B, BG and G refer to butyrate, butyrate with glucose and glucose feeding. The right axis is for the controls while the left axis is for the enriched conditions B, BG, and G.

When glucose was the substrate, no production of CH4 was observed after stopping completely H2:CO2 (∼6 days) and acetate (∼20 days) feedings (**Figure 1**). However, more than 95% of the glucose was consumed after 10 days, while H2, ethanol, acetate, butyrate, propionate and caproate were produced (**Supplementary Figure 1**; **Supplementary data 1**). This indicates a metabolism oriented towards acidogenesis and chain elongation rather than methanogenesis. Conditions supplied with butyrate and glucose did not produce CH4 overall. Only very small amounts were detected in P-BG (2-4% yields) and in M-BG-R2 (6% yield) during the last generation (**Figure 1b-c**; **Supplementary data 1**). The glucose was consumed at 90% on average after 10 days, but butyrate concentrations remained stable or increased slightly, indicating some production. Production of H2, ethanol, acetate, propionate (in P only) and caproate (in A only) was observed (**Supplementary Figure 1**). This denotes, as with glucose-fed conditions, a metabolism oriented towards acidogenesis. Finally, under conditions supplied with butyrate, a small production of CH4 was observed (1-2% yields). The A-B-R1 culture was an exception, with yields in the range of 22% to 48% during the last generation, indicating a sudden improvement in CH4 production. In this same culture, butyrate consumption ranged from 58% to 93% over the last generation, whereas it did not exceed 7% under the other butyrate-fed conditions (**Figure 1**; **Supplementary data 1**).

### Influence of substrate on microbial composition over time

Microbial diversity and composition across all conditions were monitored to assess the influence of carbon sources, amino acids, and vitamins on the consortia. According to the NMDS analysis on the 642 retained ASVs (**Figure 2c**), samples were clustered by condition: first by substrate (NMDS1 axis) with the controls and butyrate clustering together; second by inoculum (NMDS2 axis) with M and P inocula related samples on one side.

**Figure 2:**
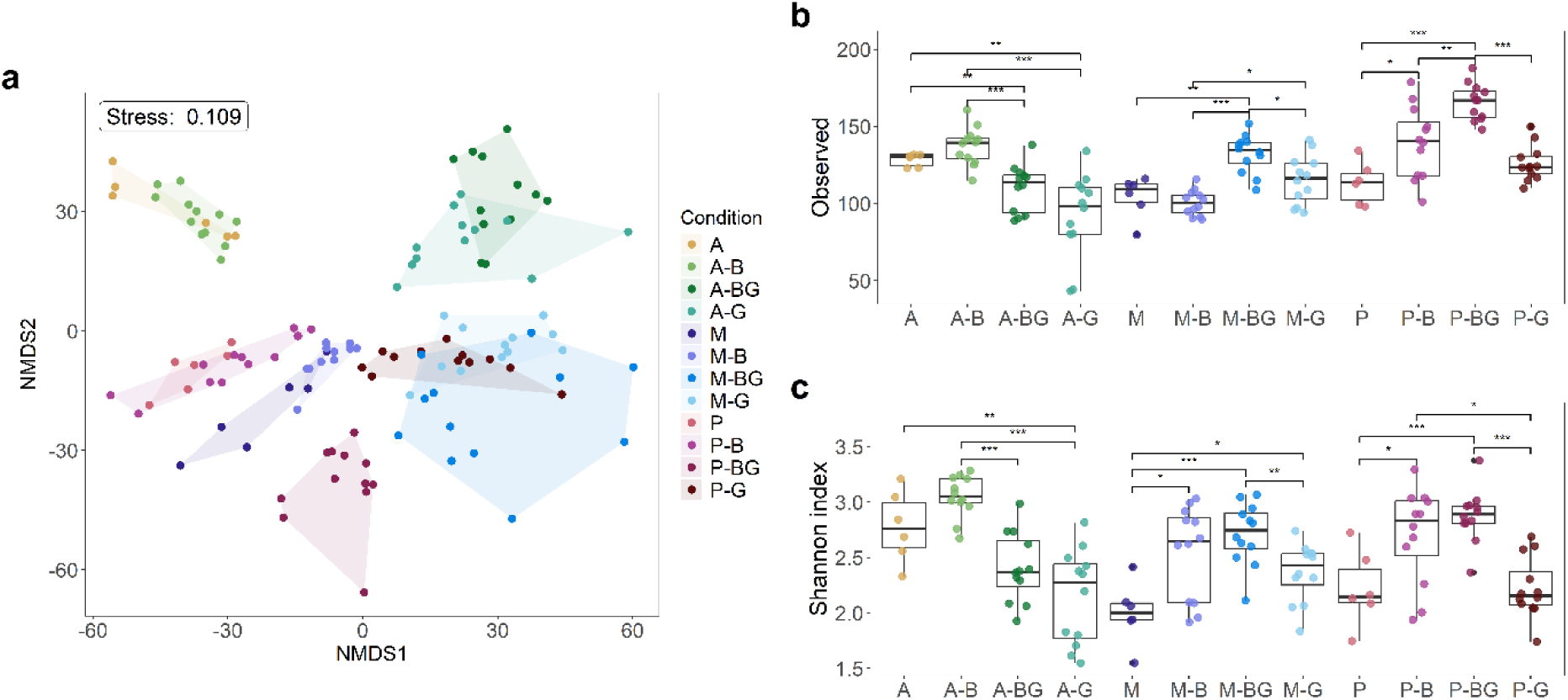
Microbial diversity indexes and repartitions between samples. **a)** Non-metric Multidimensional Scaling (NMDS) analysis for each sample at each time interval, colored by condition. **b)** Observed richness in samples at all time slots (T1-D15 to T6-D88) grouped by condition (inoculum + feeding). **c)** Shannon indexes for each sample grouped by condition. Significance brackets were calculated using ‘ggsignif’ R package: ‘***’=p-value ≤ 0.001, ‘**’≤0.01, ‘*’≤0.05. Statistics were calculated only between groups from the same inoculum. *NS’ results are not shown. A, M and P refer to inocula. B, BG, and G refer to butyrate, butyrate with glucose, and glucose feeding.

Concerning microbial diversity and composition in the controls, the A ctrl, M ctrl, and P ctrl harbored richness of 129, 105 and 113 ASVs on average from T1 (Day 15) to T6 (Day 88) (**Figure 2a**; **Supplementary Figure 2**; **Supplementary data 3**). The three controls were mainly composed of Archaea representing up to 56%, 83% and 78% of the total community at T6, respectively. In A ctrl, *Methanosarcina* was the dominant Archaea, while *Methanobacterium* and *Methanosarcina* were the dominant ones in M ctrl and P ctrl (**Figure 3**; **Supplementary data 7**). Overall, differences between A inoculum and the M and P inocula can be explained by the previous enrichment of these communities with acetate (A) or H2:CO2 (P and M). No direct effects related to conductive material were observed in this study. This may be explained by the fact that the changes induced by conductive materials on the microbial communities had already occurred during the previous enrichment period of (Giangeri, Tsapekos *et al*. 2025), leading to different communities in the inocula.

**Figure 3:**
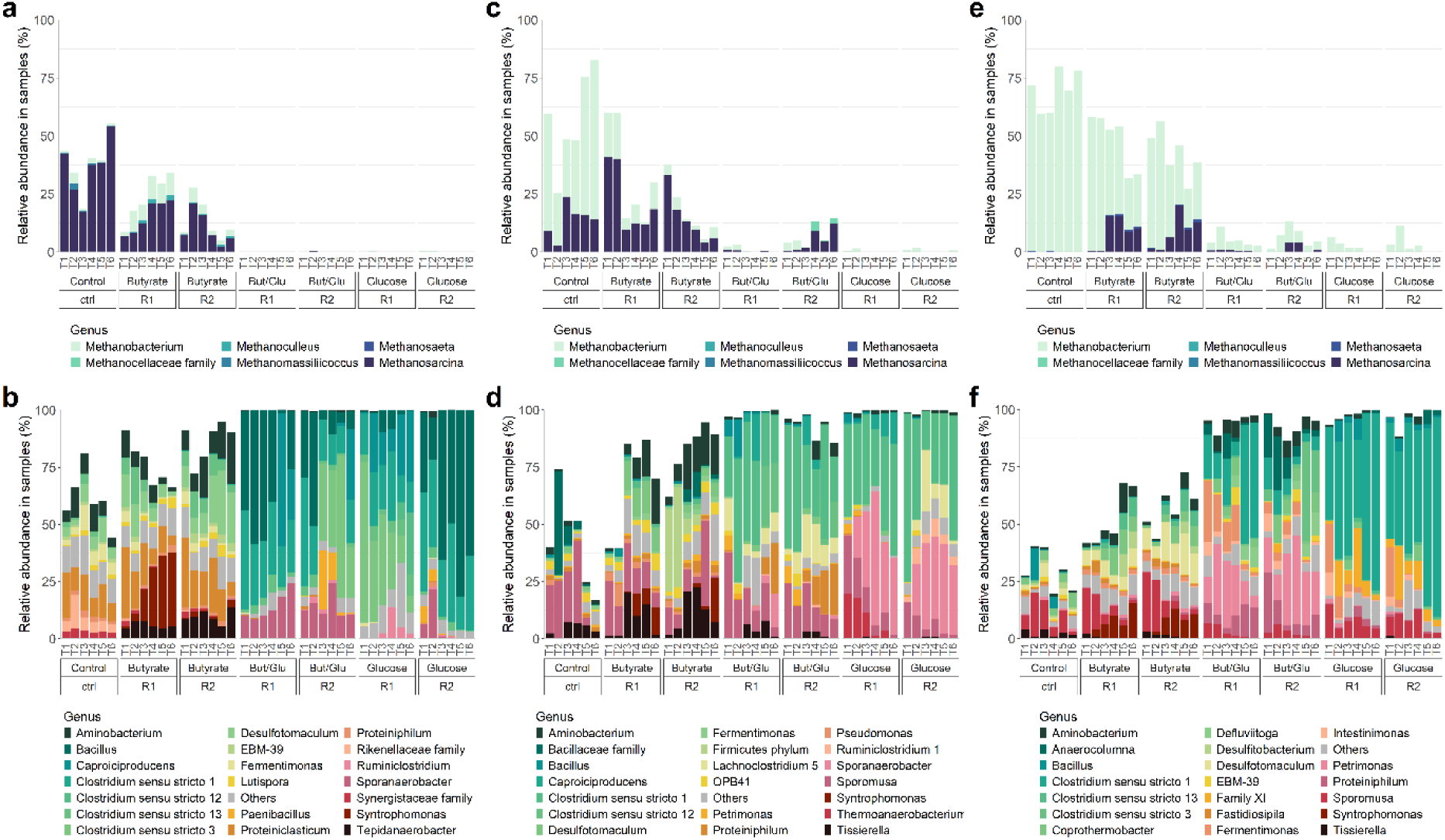
Microbial relative abundance at the genus level in enrichments and controls. All Archaeal genera are represented for **a)** A, **c)** M, and **e)** P inocula; while only the 20 most abundant Bacterial genera, calculated overall per inoculum, are represented for **b)** A, **d)** M and **f)** P inocula. The 100% scale of the y-axis considers both Archaea and Bacteria. Time intervals: T1=14 days, T2=28 days, T3=43 days, T4=57 days, T5=71 days, T6=88 days.

Regarding butyrate-fed conditions, the observed richness was similar to the controls, except for the P inoculum, where it was significantly higher. The Shannon indexes were overall substantially higher than the controls and glucose-fed conditions, indicating a higher diversity and/or abundance balance of the butyrate-fed communities (**Figure 2a-b**; **Supplementary data 3**). In all butyrate-fed conditions, Archaea relative abundance was decreasing over time, except in A-B-R1 where Archaea proportion increased up to 34% at T6, reaching proportions closer to the A ctrl (**Figure 3c-f**; **Supplementary data 7**). The main genera overall were *Methanosarcina, Aminobacterium*, *Desulfotomaculum* and *Syntrophomonas*. More particularly, *Methanobacterium* was highly abundant in P-B conditions; *Tissierella* was abundant in M-B conditions; and *Proteiniclasticum* (as in A ctrl) and *Tepidanaerobacter* were abundant in A-B communities. Only in the case of A-B-R1, a significant increase in the proportion of *Syntrophomonas* in the community was observed, rising from 1% to 33% over three months. *Methanosarcina* proportion also increased from 7% to 22% in this same culture. It is notable that when the substrate was shifted from acetate to butyrate (inoculum A) or from H2:CO2 to butyrate (inocula M and P), the bacterial communities remained quite stable, showing only minor changes. This can be explained by a balanced butyrate degradation providing both CO2 and acetate which were the original substrate of the inocula communities. Also, acetoclastic and hydrogenotrophic methanogens were maintained in all butyrate-fed conditions even though their total abundance decreased slightly and CH4 production was very low compared to the controls.

Considering butyrate/glucose feeding, the Shannon indexes were significantly lower than for the butyrate feeding, while similar to glucose feeding for the A inoculum. On the other hand, Shannon indexes were similar to butyrate feeding and significantly higher than glucose feeding in M and P inocula (**Figure 2a-b**; **Supplementary data 3**). In all butyrate/glucose-fed conditions, Archaea were subdominant (up to 0.4%) and disappeared from the community (**Figure 3**; **Supplementary data 7**). Overall, the main genera were Clostridium sensu stricto 1, *Bacillus*, *Sporanaerobacter, Caproiciproducens, Defluviitoga* (in P-BG) and *Proteiniphilum* (in M-BG and P-BG). The only exception was M-BG-R2 where the Archaea proportion increased from 4% to 15%. In this condition, *Proteiniphilum* was enriched at T6, as well as *Methanosarcina* reaching 22% and 12% of the total community, respectively.

Finally, for glucose-fed conditions, the Shannon indexes were the lowest, indicating a lower richness and/or a bigger imbalance in abundance among species compared to butyrate-fed and/or butyrate/glucose-fed conditions (**Figure 2a-b**; **Supplementary data 3**). When glucose was introduced in the cultures, the balance of the microbial community was disrupted in less than 15 days, harboring few overabundant genera (2-3 genera at >20% of relative abundance). In all glucose-fed conditions, Bacteria were dominant at T6 with almost no Archaea left (up to 1.7%), as in butyrate/glucose-fed conditions (**Figure 3**; **Supplementary data 7**). After three months, *Caproiciproducens*, *Bacillus*, Clostridium sensu stricto 1 and *Sporanaerobacter* were the most abundant genera.

Overall, after the first 15 days of exposure to a new substrate, specific functional groups were enriched, and communities remained relatively stable for three months. The main methanogen of interest, *Methanosarcina*, was maintained or enriched under butyrate-fed conditions, even though its activity was very low, while the genus almost completely disappeared when glucose was provided.

### Selection of vitamins and amino acids to support *Methanosarcina* and its potential identified partners growth during the substrate disturbance

The MAGs from (Giangeri, Tsapekos *et al*. 2025) were screened to identify potential helpers that can supply at least partly the auxotrophies of *Methanosarcina*. Analysis of KEGG modules from the three main MAGs affiliated to *Methanosarcina* in the inocula (*M. spelaei* DTU20, *M. mazei* DTU80, *M.* sp002499445 DTU142) predicted auxotrophies for the following vitamins and coenzymes: biotin, riboflavin, menaquinone, ubiquinone, lipoic acid, L-threo-Tetrahydrobiopterin, pantothenate, tetrahydrobiopterin, tetrahydrofolate, tocopherol/tocotorienol, pimeloyl-ACP, and coenzyme A. The *Methanosarcina* MAGs were also missing genes for tyrosine, phenylalanine, leucine, methionine and lysine biosynthesis. DTU142 also exhibited auxotrophies for serine and threonine biosynthesis. Based on these predicted auxotrophies for vitamins and amino acids, three MAGs were selected to support *Methanosarcina*: *Advenella* sp. DTU124, Eubacteriales sp. DTU119, and *Negativicutes sp*. DTU94 (**Supplementary data 5**). However, the three selected potential partners were not predicted to supplement all the vitamin and amino acid auxotrophies of *Methanosarcina* (no MAGs could) and were also exhibiting auxotrophies for vitamins and amino acids that could not be supplemented by their interaction with *Methanosarcina*. Consequently, these inferred auxotrophies were supplemented by adding nutrients to the medium to promote the growth of *Methanosarcina* spp. and their potential partners (**Table 1**).

### Abundance of the potential partners of *Methanosarcina* in the community over time

The total bacterial concentrations in A, P and M related conditions were relatively stable from day 28 to 57. Bacterial concentrations were overall higher in the glucose-fed and butyrate/glucose-fed conditions with on average 1.02×10^6^±6.14×10^5^ and 1.89×10^6^±1.62×10^6^ gene copies/µL respectively, compared to butyrate-fed conditions with on average 3.55×10^5^±1.40×10^5^ gene copies/µL (**Supplementary data 8**).

Concerning the MAGs related to the three selected partners, *Advenella sp.* DTU124, Eubacteriales sp. DTU119, and *Negativicutes sp*. DTU94, no enrichment of related species was observed despite the addition of vitamins and amino acids that should have complemented their auxotrophies and promoted their growth. They remained stable in concentration and in very low abundance (<10^4^ gene copies/µL) within the community or disappeared entirely. This outcome may be explained by the extremely low abundances of the target partner species already at the beginning of the experiment and by potential lack of adequate substates supporting their growth.

### Enrichment of specific non-targeted taxa and their gene content

The microbial diversity profiling highlighted the growth of several taxa under specific conditions that were not targeted. First, the *Syntrophomonas* genus (DTU160, DTU167 and DTU169) was found enriched in each butyrate-fed conditions. The genus relative abundance started to increase after 15 days in A and P inocula related conditions, and a bit later after 43-57 days in M inoculum related conditions (**Supplementary data 7**; **Supplementary data 9**). The MAGs Syntrophomonadaceae sp. DTU160 and DTU167 exhibited genes related to butyrate oxidation: acetate CoA/acetoacetate CoA-transferase alpha subunit (*atoD*) and beta subunit (*atoA*) for the former; butyrate kinase (*buk*) for the latter (**Supplementary data 10**). The genus *Tepidanaerobacter* was enriched in A-B conditions (**Supplementary data 7**; **Supplementary data 9**). The related MAG Thermoanaerobacterales sp. DTU146 harbored auxotrophies for folic acids and phylloquinone, which were supplemented in conditions related to A inoculum.

The genus *Aminobacterium,* most likely *A. mobile* DTU9 according to relative abundances, and *Sporanaerobacter,* likely S. *acetigenes* DTU16 based on relative abundances, was enriched under butyrate-fed or glucose-fed conditions respectively, predominantly in cultures derived from inocula A and M where tyrosine and phenylalanine were supplemented (**Supplementary data 9**). Genes related to tyrosine and phenylalanine degradation pathways, such as the histidinol-phosphate aminotransferase (*hisC*; K00817) and amidase (*amiE*; K01426) were found in DTU9 and DTU163 (**Supplementary data 11**).

## Discussion

The carbon source provided was the main factor structuring the microbial community distribution across samples. As a higher VFA, butyrate plays a crucial role in acetogenesis degradation process, providing in turn substrates for methanogenesis (Xu Y *et al*. 2023, Shi X *et al*. 2025). Its degradation relies on syntrophic interactions between butyrate-oxidizing bacteria and methanogens. Butyrate is first degraded into acetate and/or CO2, which are then used by methanogens depending on their substrate affinity (Schink and Stams 2012, Nikitina *et al*. 2023). Accordingly, the microbial composition remained quite preserved under butyrate-fed conditions compared to the controls, even after three months, with an enrichment in syntrophic butyrate-oxidizing bacteria (SBOB) such as *Syntrophomonas* (McInerney *et al*. 1981), syntrophic acetate-oxidizing bacteria (SAOB) that feed on the acetate produced like *Tepidanaerobacter* (DTU146) (Westerholm, Roos, and Schnürer 2011), acidogenic and hydrogen-producer bacteria like *Coprothermobacter* (DTU70) (Gagliano *et al*. 2015), and finally some sulfate-reducing bacteria (SRB) genera such as *Tissierella* and *Desulfotomaculum* (DTU177) (Aüllo *et al*. 2013, Ikkert *et al*. 2013) using most probably the H2 produced and keeping it low. The slow growth of SBOB in most conditions may explain the substrate limitation of methanogens and the subsequent low production of methane. The study of (Ziels, Beck, and Stensel 2017) reported *Syntrophomonas* enrichment (up to ∼20% of the community) after ∼41 days in continuous and pulse-fed bioreactors, which are coherent with our observations in some of the cultures. Nevertheless, they were working on more diverse microbial communities from feedstock, while in our case a loss of SBOB, and acetogenesis functional group in general, could have occurred due to the lack of higher substrates during the previous enrichment period and the subsequent strong adaptation of the inoculum-derived communities to acetate and CO2 as substrates. The redevelopment of low-abundance taxa related to this functional group therefore requires an adaptation period that was not fully achieved within three months in most of the cultures studied. On the other hand, Bacteria of the Clostridia class responsible for acidogenesis started growing rapidly when glucose was introduced as substrate, leading to rapid accumulation of VFA, an indicator of imbalance in the AD systems (Boe *et al*. 2010, Pratt *et al*. 2012). Acetogenic bacteria, responsible for degradation of VFA, were unable to oxidize VFA at the same rate as they were produced, while high H2 production may also have inhibited syntrophic acetogenic interactions (Sieber, McInerney, and Gunsalus 2012). The rapid and efficient resumption of growth in Clostridia species was predictable and can be explained by the ability of this taxonomic group to form spores and remain dormant for long periods before re-emerging when conditions become more favorable, e.g., new substrate availability (Labbé and Remi Shih 1997, Olguín-Araneda *et al*. 2015).

Overall, promoting acetogenesis through butyrate feeding helped to maintain a balance between syntrophic bacteria and methanogens and promoted the re-diversification of the simplified community, while glucose enriched acidogenic bacteria, disrupting the community balance and led at the end to less diversify microbial communities. This highlights the importance of going steps by steps in reintroducing functional groups in a simplified community oriented towards methanogenesis, as a well-structured acetogenic community is first needed to be able to handle the VFA produced upstream. Moreover, variability could be observed between replicate regarding the growth resumption of some taxa and can be explain by the addition of several complex and hardly predictable factors such as the dilution of the low-abundance taxa in the inoculum, mortality due to stress, and the spatial arrangement of the microbial community (Abreu *et al*. 2019, Micali *et al*. 2023).

The maintenance of methanogenesis with the change of substrate was achieved only in few conditions fed with butyrate. The supplementation in vitamins and amino acids didn’t directly help the growth or maintenance of *Methanosarcina* and its potential microbial partners in the communities, but the functional redundancy of the communities has led to the enrichment of microorganisms for the same function and helped *Methanosarcina* in several specific cases.

These targeted microbial partners may not have exhibited the most suitable or competitive metabolism to the new induced condition, compared to the other enriched species. The amino acids supplementation did promote the growth of taxonomic groups, such as members of the *Aminobacterium* genus which are known as amino acid degraders, especially when co-culture with hydrogenotrophs, such as *Methanobacterium* (Baena *et al*. 2000, Shi Z *et al*. 2021). This genus was consistently present and enriched in the two methane-producing cultures and has been positively correlated in the literature as a potential syntrophic partner of *Methanosarcina* (Yin *et al*. 2025). Vitamins and folic acids helped to enrich taxa such as *Tepidanaerobacter*, which could have benefited from the supplements in folic acids and phylloquinone in inoculum A. This genus has been documented as syntrophic acetate-oxidizers when co-cultivated with hydrogenotrophic methanogens and could have also benefited from their syntrophic relationship with the enriched *Methanosarcina* sp002499445 DTU142 and *Methanobacterium* sp. DTU30, thereby providing them in turn with H2 and CO2 (Westerholm, Roos, and Schnürer 2011). As a final note, vitamin and amino acid auxotrophies did not appear to be as limiting as expected in the medium (they did not induce many species enrichment), which may be explained by the already established balance between low-abundance microorganisms producing the essential nutrients needed by the community and the beneficiaries, which are not necessarily in direct interaction (Morris, Lenski, and Zinser 2012). The fact that the three inocula exposed to similar substrates but variable amino acids and vitamins end up with similar functional groups highlights a strong functional convergence rather than a taxonomical convergence and the interchangeability of species for similar functions.

To conclude, the approach used in this study allowed us to successfully re-diversify the simplified methanogenic microbial community when targeting the acetogenic functional group, even after months of methanogenic substrate feeding, demonstrating the resilience of such community and the importance of low-abundance taxa. The results also highlighted the need for transition steps in the complexification of the substrate, with the need to favorize first acetogenesis, then acidogenesis, to avoid VFA accumulation. The balance of the AD system and the production of methane was difficult to maintain through this period of disturbance and can be explained by substrate overload and the time needed for the higher functional groups to re-develop. The second objective of using predicted auxotrophies for vitamin and amino acid to support and maintain the growth of *Methanosarcina* and targeted microbial partners during the substrate disturbance highlighted the need to target function groups rather than specific taxa to leverage the high functional redundancy of AD consortia. Together, these results shed new light on the key factors to consider in the long-term perspective of designing simplified communities and tailored-made microbial consortia for AD. To go further, it would be interesting to investigate whether within the same community, after several cycles of alternating between substrates of low and high complexity, the same species reappear, or whether a new microbial rearrangement is favored each time, with functional convergence being the sole determining factor.

## Availability of data and materials

The datasets supporting the conclusions of this article are included within the article and its additional files. Scripts for processing sequencing data and generating figures are available in the GitHub repository rpeguil/MethanoEnrich. The raw sequencing data and MAGs for this study have been deposited in the European Nucleotide Archive (ENA) at EMBL-EBI under accession number PRJEB98414 (https://www.ebi.ac.uk/ena/browser/view/PRJEB98414), samples SAMEA120801592 to SAMEA120801598 for shotgun metagenomics, samples SAMEA120799023 to SAMEA120799148 for amplicon sequencing and analysis accession GCA_982384175 to GCA_982388805 for MAGs. The sequences of the MAGs are also available as an archive in the DTU collection repository, https://doi.org/10.11583/DTU.c.8263735. For the A inoculum, raw data were deposited under NCBI BioProject PRJNA1221661, sample SAMN46749652. Data from the study of (Giangeri, Tsapekos *et al*. 2025) were used and can be found under the NCBI BioProject PRJNA1063026.

## Acknowledgments

The authors would like to thank the sequencing platform of Technical University of Denmark (DTU) National Food Institute for the shotgun sequencing of the two P and M inocula. The authors would also like to thank Dr. Margot Mahieux of DTU for her advice, which helped improve the manuscript, as well as Dr. Antonio Grimalt Alemany of DTU for the graphic representation using BioRender. Reference for the featured image: created in BioRender.

Grimalt, A. (2027) https://BioRender.com/w1c3kic.

## Authors’ contributions

RP contributed to the study design, data acquisition, analysis, and interpretation, as well as to the writing of the first draft and final manuscript. GG made a substantial contribution to the study conception and thoroughly reviewed the manuscript. MMJ contributed to data acquisition and interpretation, as well as manuscript revision. IA contributed substantially to the conceptualization of the work and extensively revised the manuscript.

## References

Abel S, Abel zur Wiesch P, Davis BM et al. Analysis of Bottlenecks in Experimental Models of Infection. PLoS Pathog 2015;11(6):e1004823. 10.1371/JOURNAL.PPAT.1004823.

Abreu CI, Friedman J, Andersen Woltz VL et al. Mortality causes universal changes in microbial community composition. Nature Communications 2019 10:1 2019;10(1):2120-. 10.1038/s41467-019-09925-0.

Adekunle KF, Okolie JA. A Review of Biochemical Process of Anaerobic Digestion. Advances in Bioscience and Biotechnology 2015;06(03):205–12. 10.4236/ABB.2015.63020.

Agler MT, Spirito CM, Usack JG et al. Development of a highly specific and productive process for n-caproic acid production: applying lessons from methanogenic microbiomes. Water Science and Technology 2014;69(1):62–8. 10.2166/WST.2013.549.

Alneberg J, Bjarnason BS, Bruijn I De et al. Binning metagenomic contigs by coverage and composition. Nature Methods 2014 11:11 2014;11(11):1144–6. 10.1038/nmeth.3103.

Andrews S. FastQC: a quality control tool for high throughput sequence data. Preprint, Babraham Bioinformatics, 2010.

Angelidaki I, Sanders W. Assessment of the anaerobic biodegradability of macropollutants. Rev Environ Sci Biotechnol 2004;3(2):117–29. 10.1007/S11157-004-2502-3/METRICS.

Apprill A, Mcnally S, Parsons R et al. Minor revision to V4 region SSU rRNA 806R gene primer greatly increases detection of SAR11 bacterioplankton. Aquatic Microbial Ecology 2015;75(2):129–37. 10.3354/ame01753.

Aroney STN, Newell RJP, Nissen JN et al. CoverM: read alignment statistics for metagenomics. Bioinformatics 2025;41(4). 10.1093/BIOINFORMATICS/BTAF147.

Aüllo T, Ranchou-Peyruse A, Ollivier B et al. Desulfotomaculum spp. and related gram-positive sulfate-reducing bacteria in deep subsurface environments. Front Microbiol 2013;4(DEC):58161. 10.3389/FMICB.2013.00362/FULL.

Baena S, Fardeau ML, Labat M et al. Aminobacterium mobile sp. nov., a new anaerobic amino-acid-degrading bacterium. In: International Journal of Systematic and Evolutionary Microbiology, vol. 50. 2000.

Bernard M, RuCrossed D sign© O, Mariadassou M et al. FROGS: a powerful tool to analyse the diversity of fungi with special management of internal transcribed spacers. Brief Bioinform 2021;22(6). 10.1093/bib/bbab318.

Boe K, Batstone DJ, Steyer JP et al. State indicators for monitoring the anaerobic digestion process. Water Res 2010;44(20):5973–80. 10.1016/J.WATRES.2010.07.043.

Bolger AM, Lohse M, Usadel B. Trimmomatic: A flexible trimmer for Illumina sequence data. Bioinformatics 2014;30(15):2114–20. 10.1093/bioinformatics/btu170.

Callahan BJ, McMurdie PJ, Rosen MJ et al. DADA2: High-resolution sample inference from Illumina amplicon data. Nature Methods 2016 13:7 2016;13(7):581–3. 10.1038/nmeth.3869.

Chang H, Yin Q, He K et al. Feeding regime selectively enriching acetoclastic methanogens to enhance energy production in anaerobic digestion systems. Biochem Eng J 2025;220:109764. 10.1016/J.BEJ.2025.109764.

Chaumeil PA, Mussig AJ, Hugenholtz P et al. GTDB-Tk v2: memory friendly classification with the genome taxonomy database. Bioinformatics 2022;38(23):5315–6. 10.1093/BIOINFORMATICS/BTAC672.

Chen Y, Cheng JJ, Creamer KS. Inhibition of anaerobic digestion process: A review. Bioresour Technol 2008;99(10):4044–64. 10.1016/j.biortech.2007.01.057.

Eren AM, Kiefl E, Shaiber A et al. Community-led, integrated, reproducible multi-omics with anvi’o. Nature Microbiology 2020 6:1 2020;6(1):3–6. 10.1038/s41564-020-00834-3.

Escudié F, Auer L, Bernard M et al. FROGS: Find, Rapidly, OTUs with Galaxy Solution. Bioinformatics 2018;34(December 2017):1287–94. 10.1093/bioinformatics/btx791.

Gagliano MC, Braguglia CM, Petruccioli M et al. Ecology and biotechnological potential of the thermophilic fermentative Coprothermobacter spp. FEMS Microbiol Ecol 2015;91(5):18. 10.1093/FEMSEC/FIV018.

Galperin MY, Vera Alvarez R, Karamycheva S et al. COG database update 2024. Nucleic Acids Res 2025;53(D1):D356–63. 10.1093/NAR/GKAE983,.

Giangeri G, Campanaro S, Kyrpides NC et al. Unlocking the potential of designed microbial consortia: A breakthrough for sustainable waste management and climate resilience. Environmental Science and Ecotechnology 2025;25:100558. 10.1016/J.ESE.2025.100558.

Giangeri G, Tsapekos P, Pitsikoglou D et al. Deciphering direct interspecies electron transfer activity in microbial interactions: The influence of conductive materials in anoxic ecosystems. Chemical Engineering Journal 2025;503:158716. 10.1016/J.CEJ.2024.158716.

Gilmore SP, Lankiewicz TS, Wilken SE et al. Top-Down Enrichment Guides in Formation of Synthetic Microbial Consortia for Biomass Degradation. ACS Synth Biol 2019;8(9):2174–85. 10.1021/ACSSYNBIO.9B00271.

Gloor GB, Macklaim JM, Pawlowsky-Glahn V et al. Microbiome datasets are compositional: And this is not optional. In: Frontiers in Microbiology, no. NOV. Preprint, 2017, 8.1–6. 10.3389/fmicb.2017.02224.

Goonesekera EM, Grimalt-Alemany A, Thanasoula E et al. Biofilm mass transfer and thermodynamic constraints shape biofilm in trickle bed reactor syngas biomethanation. Chemical Engineering Journal 2024;500:156629. 10.1016/J.CEJ.2024.156629.

Hu H, Wang M, Huang Y et al. Guided by the principles of microbiome engineering: Accomplishments and perspectives for environmental use. MLife 2022;1(4):382–98. 10.1002/MLF2.12043;JOURNAL:JOURNAL:2770100X;WGROUP:STRING:PUBLICATION.

Hubalek V, Buck M, Tan B et al. Vitamin and Amino Acid Auxotrophy in Anaerobic Consortia Operating under Methanogenic Conditions. MSystems 2017;2(5). 10.1128/MSYSTEMS.00038-17/ASSET/06F74525-9172-4DA6-B16F-DC5A841FA097/ASSETS/GRAPHIC/SYS0051721440003.JPEG.

Ikkert OP, Gerasimchuk AL, Bukhtiyarova PA et al. Characterization of precipitates formed by H2S-producing, cu-resistant firmicute isolates of tissierella from human gut and desulfosporosinus from mine waste. *Antonie van Leeuwenhoek*, International Journal of General and Molecular Microbiology 2013;103(6):1221–34. 10.1007/S10482-013-9900-X/FIGURES/8.

J Cira N, Pearce MT, Quake SR. Neutral and selective dynamics in a synthetic microbial community. Proc Natl Acad Sci U S A 2018;115(42):E9842–8. 10.1073/PNAS.1808118115;PAGE:STRING:ARTICLE/CHAPTER.

Jourdain L, Gu W. Designing synthetic microbial communities for enhanced anaerobic waste treatment. In: Applied and Environmental Microbiology, vol. 91, no. 6. Preprint, American Society for Microbiology, 1 Jun. 2025. 10.1128/aem.00404-25.

Kanehisa M, Furumichi M, Sato Y et al. KEGG: biological systems database as a model of the real world. Nucleic Acids Res 2025;53(D1):D672–7. 10.1093/NAR/GKAE909.

Kanehisa M, Furumichi M, Tanabe M et al. KEGG: new perspectives on genomes, pathways, diseases and drugs. Nucleic Acids Res 2017;45(D1):D353–61. 10.1093/NAR/GKW1092.

Kanehisa M, Goto S. KEGG: Kyoto Encyclopedia of Genes and Genomes. Nucleic Acids Res 2000;28(1):27–30. 10.1093/NAR/28.1.27.

Kang DD, Froula J, Egan R et al. MetaBAT, an efficient tool for accurately reconstructing single genomes from complex microbial communities. PeerJ 2015;2015(8):e1165. 10.7717/PEERJ.1165/SUPP-2.

Kang DD, Li F, Kirton E et al. MetaBAT 2: An adaptive binning algorithm for robust and efficient genome reconstruction from metagenome assemblies. PeerJ 2019;2019(7):e7359. 10.7717/PEERJ.7359/SUPP-3.

Labbé RG, Remi Shih NJ. Physiology of Sporulation of Clostridia. The Clostridia 1 Jan. 1997:21–32. 10.1016/B978-012595020-6/50004-8.

Langmead B, Salzberg SL. Fast gapped-read alignment with Bowtie 2. Nat Methods 2012;9(4):357–9. 10.1038/nmeth.1923.

Li D, Liu CM, Luo R et al. MEGAHIT: An ultra-fast single-node solution for large and complex metagenomics assembly via succinct de Bruijn graph. Bioinformatics 2015;31(10):1674–6. 10.1093/bioinformatics/btv033.

Lyu X, Nuhu M, Candry P et al. Top-down and bottom-up microbiome engineering approaches to enable biomanufacturing from waste biomass. J Ind Microbiol Biotechnol 2024;51(2). 10.1093/JIMB/KUAE025.

McInerney MJ, Bryant MP, Hespell RB et al. Syntrophomonas wolfei gen. nov. sp. nov., an Anaerobic, Syntrophic, Fatty Acid-Oxidizing Bacterium. Appl Environ Microbiol 1981;41(4):1029–39. 10.1128/AEM.41.4.1029-1039.1981.

Micali G, Hockenberry AM, Co AD et al. Minorities drive growth resumption in cross-feeding microbial communities. Proc Natl Acad Sci U S A 2023;120(45):e2301398120. 10.1073/PNAS.2301398120;PAGE:STRING:ARTICLE/CHAPTER.

Morris JJ, Lenski RE, Zinser ER. The Black Queen Hypothesis: Evolution of Dependencies through Adaptive Gene Loss. MBio 2012;3(2). 10.1128/mBio.00036-12.

Nikitina AA, Kallistova AY, Grouzdev DS et al. Syntrophic Butyrate-Oxidizing Consortium Mitigates Acetate Inhibition through a Shift from Acetoclastic to Hydrogenotrophic Methanogenesis and Alleviates VFA Stress in Thermophilic Anaerobic Digestion. Applied Sciences (Switzerland) 2023;13(1). 10.3390/app13010173.

Olguín-Araneda V, Banawas S, Sarker MR et al. Recent advances in germination of Clostridium spores. Res Microbiol 2015;166(4):236–43. 10.1016/J.RESMIC.2014.07.017.

Olm MR, Brown CT, Brooks B et al. dRep: a tool for fast and accurate genomic comparisons that enables improved genome recovery from metagenomes through de-replication. ISME J 2017;11(12):2864–8. 10.1038/ISMEJ.2017.126.

Parada AE, Needham DM, Fuhrman JA. Every base matters: Assessing small subunit rRNA primers for marine microbiomes with mock communities, time series and global field samples. Environ Microbiol 2016;18(5):1403–14. 10.1111/1462-2920.13023.

Parks DH, Chuvochina M, Rinke C et al. GTDB: an ongoing census of bacterial and archaeal diversity through a phylogenetically consistent, rank normalized and complete genome-based taxonomy. Nucleic Acids Res 2022;50(D1):D785–94. 10.1093/NAR/GKAB776.

Pratt S, Liew D, Batstone DJ et al. Inhibition by fatty acids during fermentation of pre-treated waste activated sludge. J Biotechnol 2012;159(1–2):38–43. 10.1016/J.JBIOTEC.2012.02.001.

Price MN, Deutschbauer AM, Arkin AP. GapMind: Automated Annotation of Amino Acid Biosynthesis. MSystems 2020;5(3). 10.1128/MSYSTEMS.00291-20/SUPPL_FILE/REVIEWER-COMMENTS.PDF.

Quast C, Pruesse E, Yilmaz P et al. The SILVA ribosomal RNA gene database project: Improved data processing and web-based tools. Nucleic Acids Res 2013;41(D1):590–6. 10.1093/nar/gks1219.

R Core Team (2019). R Core Team (2019). R: A language and environment for statistical computing. Foundation for Statistical Computing, Vienna, Austria. n.d. http://www.r-project.org/index.html (13 Apr. 2020, date last accessed).

Schink B, Stams AJM. Syntrophism among prokaryotes. The Prokaryotes: Prokaryotic Communities and Ecophysiology 1 Apr. 2012:471–93. 10.1007/978-3-642-30123-0_59/FIGURES/00594.

Shade A, Jones SE, Gregory Caporaso J et al. Conditionally rare taxa disproportionately contribute to temporal changes in microbial diversity. MBio 2014;5(4):1371–85. 10.1128/MBIO.01371-14/FORMAT/EPUB.

Shi X, Yasuda S, Wang Z et al. Microbial transitions and degradation pathways driven by butyrate concentration in mesophilic and thermophilic anaerobic digestion under low hydrogen partial pressure. Bioresour Technol 2025;419:132012. 10.1016/J.BIORTECH.2024.132012.

Shi Z, Campanaro S, Usman M et al. Genome-Centric Metatranscriptomics Analysis Reveals the Role of Hydrochar in Anaerobic Digestion of Waste Activated Sludge. Environ Sci Technol 2021;55(12):8351–61. 10.1021/ACS.EST.1C01995.

Sieber JR, McInerney MJ, Gunsalus RP. Genomic insights into syntrophy: The paradigm for anaerobic metabolic cooperation. Annu Rev Microbiol 2012;66:429–52. 10.1146/annurev-micro-090110-102844.

Tukanghan W, Hupfauf S, Gómez-Brandón M et al. Symbiotic Bacteroides and Clostridium-rich methanogenic consortium enhanced biogas production of high-solid anaerobic digestion systems. Bioresour Technol Rep 2021;14:100685. 10.1016/J.BITEB.2021.100685.

Uritskiy G V., Diruggiero J, Taylor J. MetaWRAP—a flexible pipeline for genome-resolved metagenomic data analysis. Microbiome 2018;6(158):1–13. 10.1186/S40168-018-0541-1.

Vayena G, Ghofrani-Isfahani P, Ziomas A et al. Impact of biochar on anaerobic digestion process and microbiome composition; focusing on pyrolysis conditions for biochar formation. Renew Energy 2024;237. 10.1016/j.renene.2024.121569.

Veseli I, Chen YT, Schechter MS et al. Microbes with higher metabolic independence are enriched in human gut microbiomes under stress. Elife 2025;12. 10.7554/ELIFE.89862.

Vrieze J De, Hennebel T, Boon N et al. Methanosarcina: The rediscovered methanogen for heavy duty biomethanation. In: Bioresource Technology. Preprint, May 2012, 112.1–9. 10.1016/j.biortech.2012.02.079.

Westerholm M, Roos S, Schnürer A. Tepidanaerobacter acetatoxydans sp. nov., an anaerobic, syntrophic acetate-oxidizing bacterium isolated from two ammonium-enriched mesophilic methanogenic processes. Syst Appl Microbiol 2011;34(4):260–6. 10.1016/J.SYAPM.2010.11.018.

Wu YW, Simmons BA, Singer SW. MaxBin 2.0: an automated binning algorithm to recover genomes from multiple metagenomic datasets. Bioinformatics 2016;32(4):605–7. 10.1093/BIOINFORMATICS/BTV638.

Xu F, Li Y, Ge X et al. Anaerobic digestion of food waste – Challenges and opportunities. Bioresour Technol 2018;247:1047–58. 10.1016/J.BIORTECH.2017.09.020.

Xu Y, Meng X, Song Y et al. Effects of different concentrations of butyrate on microbial community construction and metabolic pathways in anaerobic digestion. Bioresour Technol 2023;377:128845. 10.1016/J.BIORTECH.2023.128845.

Yin Q, Liu C, Li B et al. Expanding methanogens with genetic potential for extracellular electron transfer capabilities in anaerobic wastewater treatment ecosystems. Nature Water 2025 8 Oct. 2025:1–13. 10.1038/S44221-025-00524-6.

Ziels RM, Beck DAC, Stensel HD. Long-chain fatty acid feeding frequency in anaerobic codigestion impacts syntrophic community structure and biokinetics. Water Res 2017;117(6):218–29. 10.1016/j.watres.2017.03.060.

