## Supplementary Materials and Methods and Figures for "Increased substrate complexity drives re-diversification and functional reorganization in simplified methanogenic consortia"

#### MAGs refinement

The three MAGs affiliated with *Methanosarcina* genus were refined manually to ensure minimal contamination of these MAGs for further analyses. The Anvi'o 8 platform [36] was used with the program *anvi-run-workflow* [76], option 'metagenomics' workflow, using the raw reads from NCBI BioProject PRJNA1063026 [7] and the program *anvi-refine* [77], following the online tutorial https://merenlab.org/2015/05/11/anvi-refine/.
*Methanosarcina* MAGs statistics went from 82.9%, 94.7%, and 97.4% of completeness and 5.3%, 1.3%, and 1.3% of contamination for DTU20, DTU80 and DTU142 respectively, to 82.9%, 94.7% and 96.1% of completeness and 2.6%, 0% and 0% of contamination.

#### Composition of the vitamins solution x10

| | |
|---|---|
| - Biotin, vitamin B7 | 0.02 g/L |
| - Cyanocobalamine, vitamin B12 | 1 mg/L |
| - Pyridoxin HCL, vitamin B6 | 0.1 g/L |
| - Riboflavin, vitamin B2 | 0.05 g/L |
| - Thiamine HCL, vitamin B1 | 0.05 g/L |
| - Folic acid | 0.02 g/L |
| - Lipoic acids, thiotic acid | 0.05 g/L |
| - Nicotinic acid | 0.05 g/L |
| - DL-panthothenic acid | 0.05 g/L |
| - P-aminobenzoic acid, PABA | 0.05 g/L |

#### References

1. Giangeri G, Tsapekos P, Pitsikoglou D, Ghiotto G, Tracy Hong Lin MK, Treu L, et al. Deciphering direct interspecies electron transfer activity in microbial interactions: The influence of conductive materials in anoxic ecosystems. Chemical Engineering Journal [Internet]. Elsevier; 2025b [cited 2025 May 29];503:158716. https://doi.org/10.1016/J.CEJ.2024.158716

2. Eren AM, Kiefl E, Shaiber A, Veseli I, Miller SE, Schechter MS, et al. Community-led, integrated, reproducible multi-omics with anvi'o. Nature Microbiology 2020 6:1 [Internet]. Nature Publishing Group; 2020 [cited 2024 Feb 6];6:3–6. https://doi.org/10.1038/s41564-020-00834-3

3. Shaiber A, Willis AD, Delmont TO, Roux S, Chen LX, Schmid AC, et al. Functional and genetic markers of niche partitioning among enigmatic members of the human oral microbiome. Genome Biol [Internet]. BioMed Central Ltd; 2020 [cited 2025 May 29];21:1–35. https://doi.org/10.1186/S13059-020-02195-W;TYPE=ARTICLE;KWRD=METAGENOMICS,METAPANGENOMICS,NICHE

4. Eren AM, Esen OC, Quince C, Vineis JH, Morrison HG, Sogin ML, et al. Anvi'o: An advanced analysis and visualization platformfor 'omics data. PeerJ [Internet]. PeerJ Inc.; 2015 [cited 2025 May 29];2015:e1319. https://doi.org/10.7717/PEERJ.1319/SUPP-5

### Supplementary Figures and data

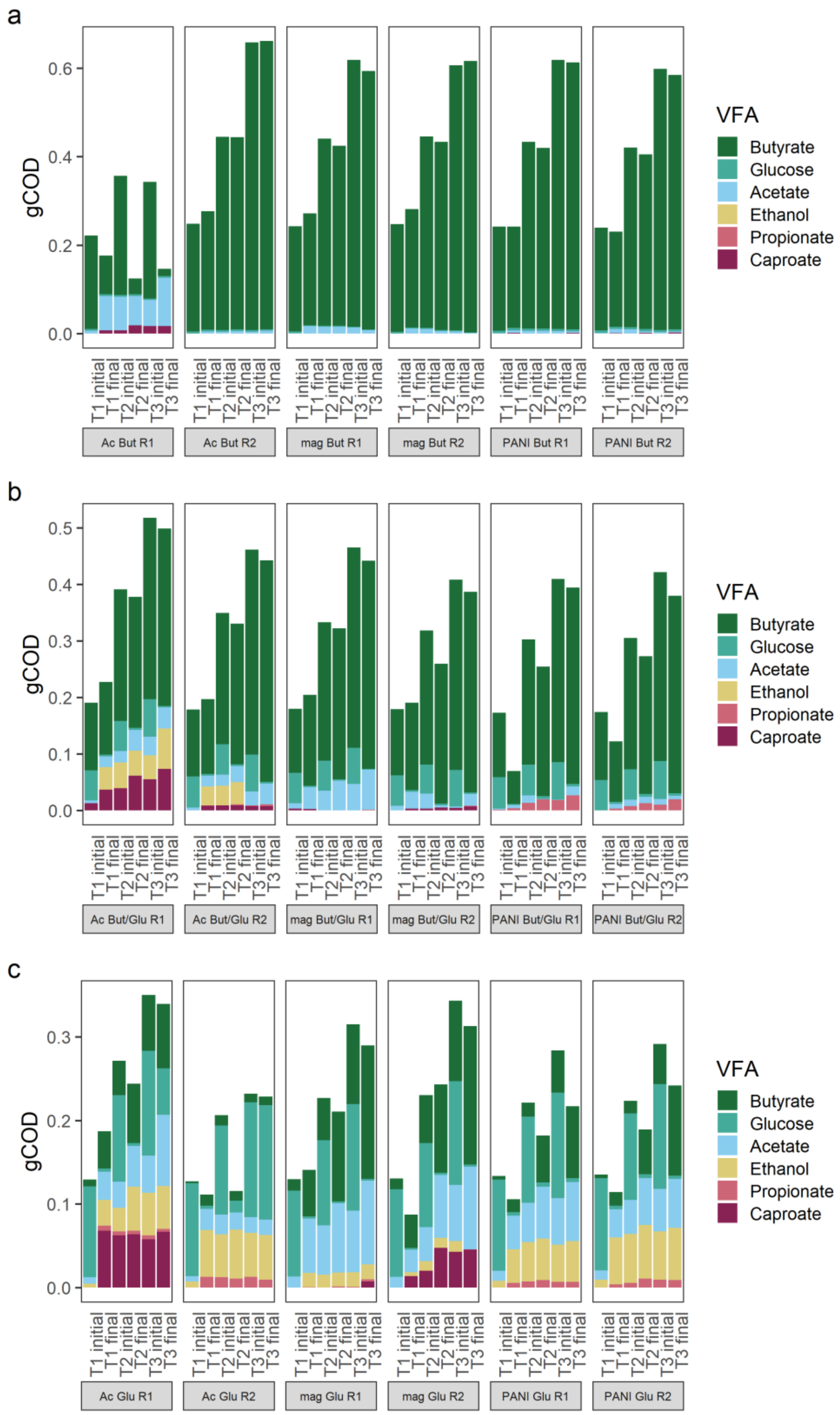


**Supplementary Figure 1:** Volatile fatty acids (VFAs) content in gCOD over the last generation of cultivations for **a)** butyrate (But), **b)** butyrate and glucose (But/Glu), and **c)** glucose (Glu) fed conditions. T1: day 67; T2: day 77; T3: day 88. Initial: refer to the sampling point just after feeding. Final: refer to sampling point after 10-11 days just before feeding. Ac, mag and PANI refer to the Acetate, $H_2:CO_2$ magnetite and $H_2:CO_2$ PANI inoculums. R1 and R2 refer to replicate 1 and 2 respectively.

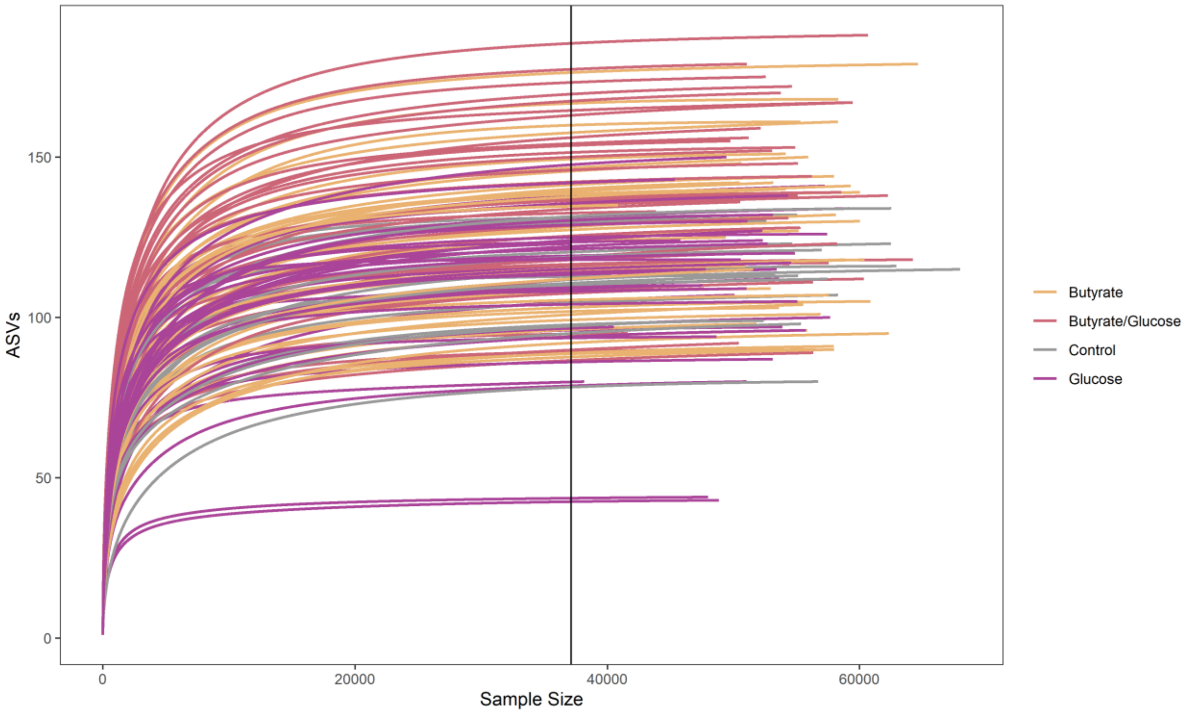


**Supplementary Figure 2**: Rarefaction curve for each metabarcoding sample, colored by feeding.

**Supplementary data 1 (.xlsx):** Data from $CH_4$ and $H_2$ measurement over the three months of enrichment, and from VFAs quantification with calculation of production yield from substrate and the carbon oxygen demand (COD) mass balance over the last month of experiment. Ctrl: refers to the 3 control cultures Ac (acetate), mag ($H_2$:$CO_2$ magnetite), and PANI ($H_2$:$CO_2$ PANI). Enrich: refers to the enriched samples with amino acids, vitamins and different subtract (But: Butyrate; Glu: Glucose).

**Supplementary data 2:** DNA concentrations per sample in ng/µL.

| [DNA] ng/µL | T1 | T2 | T3 | T4 | T5 | T6 |
|---|---|---|---|---|---|---|
| mag | 76.6 | 95 | 64.8 | 106 | 92 | 94.4 |
| mag But/Glu 1 | 42.4 | 35.6 | 22.4 | 28.4 | 57.4 | 36.6 |
| mag But/Glu 2 | 56.4 | 42.2 | 24.6 | 31.6 | 73.2 | 74.4 |
| mag But 1 | 66.4 | 43.8 | 15.3 | 16.3 | 12.1 | 18.7 |
| mag But 2 | 68.4 | 39.8 | 19.9 | 13 | 12.6 | 4.44 |
| mag Glu 1 | 73.8 | 78.4 | 26.8 | 63.6 | 84.2 | 62.8 |
| mag Glu 2 | 68.4 | 67 | 84.2 | 45.2 | 93.6 | 77.2 |
| PANI | 48.2 | 53.2 | 50.2 | 84.4 | 56.8 | 102 |
| PANI But/Glu 1 | 89.2 | 91.4 | 89.2 | 112 | 102 | 87 |
| PANI But/Glu 2 | 100 | 65.6 | 83.8 | 86.2 | 104 | 100 |
| PANI But 1 | 18.6 | 15.2 | 19.4 | 48.6 | 8.9 | 16.5 |
| PANI But 2 | 12.3 | 19.2 | 14.9 | 44.6 | 14.2 | 16.6 |
| PANI Glu 1 | 79.8 | 72 | 85.4 | 75.4 | 82 | 61.8 |
| PANI Glu 2 | 68.1 | 74.6 | 88.4 | 73.4 | 92.8 | 89.2 |
| Ac no PANI | 12.5 | 31 | 22.2 | 40.8 | 41.8 | 24.2 |
| Ac But/Glu 1 | 102 | 44.2 | 62.2 | 19.4 | 60.2 | 61.4 |
| Ac But/Glu 2 | 106 | 80.6 | 52 | 38.8 | 80.2 | 54.8 |
| Ac But 1 | 6.62 | 6.16 | 29.4 | 60.8 | 87.6 | 87 |
| Ac But 2 | 7.84 | 6.7 | 13.2 | 7.08 | 9.84 | 11.1 |
| Ac Glu 1 | 87.2 | 86.2 | 75.8 | 38.2 | 54.8 | 46.6 |
| Ac Glu 2 | 96.4 | 88.4 | 44.4 | 21 | 55 | 11.3 |

**Supplementary data 3 (.xlsx):** Data from the processing of amplicon sequencing reads, from raw data to ASVs filtering. The final number of ASV is considered the richness.

**Supplementary data 4:** Data from the processing of shotgun metagenomic (MG) reads, from raw data to MAGs coverage.

| | Mag_inoc | PANI_inoc | MG_AcBut1 T6 | MG_magBut1 T6 | MG_magButGlu2 T6 | MG_PANIBut2 T6 | MG_PANIButGlu1 T6 |
|---|---|---|---|---|---|---|---|
| Nb of raw reads (R1+R2) | 73,813,962 | 213,006,194 | 46,538,440 | 47,000,838 | 41,471,172 | 46,314,120 | 41,422,046 |
| After QC (R1+R2) | 63,076,660 | 178,853,598 | 45,603,510 | 45,897,250 | 40,556,192 | 45,371,276 | 40,551,840 |
| % removed | 14.55% | 16.03% | 2.01% | 2.35% | 2.21% | 2.04% | 2.10% |
| Nb of assembled contigs | 164,624 contigs | | | | | | |
| N50 | 15,568 bp | | | | | | |
| Min - Max bp contigs | 500 - 834,825 bp | | | | | | |
| Total bp assembled | 501,709,981,646,240 bp | | | | | | |
| % of reads used for the assembly | 96.31% | | | | | | |
| Nb of MAGs | 90 (+ 152 from Ginevra *et al.* 2025) | | | | | | |
| Nb of dereplicated MAGs | 196 | | | | | | |
| % of reads aligning on MAGs | 87.78% | 86.72% | 89.08% | 82.42% | 64.18% | 85.16% | 76.34% |

**Supplementary data 5 (.xlsx):** Auxotrophies for vitamins and amino acids detected with KEGG modules and Gapmind online tool.

**Supplementary data 6 (.xlsx):** List of qPCR primers used and their parameters for amplification.

**Supplementary data 7 (.xlsx):** ASVs taxonomy and raw counts, and relative abundances (%) for each genus and kingdom.

**Supplementary data 8 (.xlsx):** qPCR results for control and enriched samples. The average and standard deviation come from the duplicated measurements.

**Supplementary data 9 (.xlsx):** MAGs taxonomy, completeness, contamination and relative abundances (%) in inoculum and final enriched samples.

**Supplementary data 10 (.xlsx):** Count table of the genes related to butyrate the three MAGs affiliated to Syntrophomonadaceae sp. The full lists of KOfam detected in the MAGs are supplied.

**Supplementary data 11 (.xlsx):** Count table of the genes related to tyrosine and phenylalanine degradation for the MAGs of interest. The full lists of KOfam and COG id detected in each MAGs of interest are supplied.
